# Understanding organism-habitat relationships and critically evaluating reference areas is key to marine protected area assessment

**DOI:** 10.1101/2024.06.07.598001

**Authors:** Conner Jainese, Peter M. Carlson, Katelin Seeto, Lyndsey McNeill, Kelly Sivertson, Jennifer E. Caselle

## Abstract

Marine Protected Areas (MPAs) have been implemented globally as a conservation tool to improve the health and function of marine ecosystems. Research has focused on assessing MPA effectiveness, however certain habitats and communities are often avoided because they are difficult or expensive to monitor. Mesophotic (30-100m) rocky reef fish communities are a valuable commercial and recreational resource that is highly targeted but often overlooked in monitoring due to depth restricted sampling. We used two MPAs in California’s statewide protection network, along with parried reference sites, to test how protection status along with environmental conditions influenced the abundance and biomass of three highly targeted species with varying life histories and habitat preferences. Depth and habitat were strong predictors for all groups; ocean whitefish (*Caulolatilus princeps*), California sheephead (*Semicossyphus pulcher*), and targeted rockfish (*Sebastes spp*). The pattern of these effects however, differed between the species groups and the influence of protection was mixed. This work highlights how species with high habitat affinities benefit differently from protection, as a function of depth and habitat representation within the MPA/reference pair. To accurately evaluate MPAs, and the network as a whole, researchers must recognize organism-habitat relationships and incorporate them when assessing conservation efforts.

## Introduction

Marine Protected Areas (MPAs) have been implemented globally as a means of protecting marine ecosystems from anthropogenic pressures (Gleason et al. 2013; Sullivan-Stack et al. 2022; Maestro et al. 2019). The primary goal of most MPAs is to conserve and maintain marine biodiversity, with secondary benefits, such as increased catch rates outside of protection zones and economic stability for local communities, also predicted to occur (Murray et al. 1999; Sala and Giakoumi 2018). There is strong evidence that an MPA network, opposed to a single MPA, is more effective at broadly protecting a diverse suite of species across various life stages and habitats (Gaines et al. 2010; Grorud-Colvert et al. 2014). Well-designed MPA networks are able to incorporate nursery areas, account for ontogenetic habitat shifts, recognize different species-habitat associations, and protect vulnerable spawning grounds (Grüss et al. 2014; Olson et al. 2019). Monitoring all of these habitats is difficult and it is rare that all habitats within a single MPA or across an MPA network are effectively monitored. Monitoring priority is often given to those habitats more easily or cheaply accessed (shallow water reefs, grass beds, shorelines, etc.), while more challenging habitats are often data-poor (Field et al. 2007; Noble-James et al. 2023).

Mesophotic (30-100m) reefs represent one habitat that has been historically difficult to monitor and a growing body of research acknowledges our relatively incomplete understanding of these communities globally (Kahng et al. 2017; James et al. 2017; Cerrano et al. 2019). Mesophotic reefs support a diverse suite of commercially valuable species and it is estimated that a significant portion of commercial groundfish harvest falls within this depth zone (Miller et al. 2017; Jacquemont et al. 2024). As a result, these fishes are left under-surveyed and relatively data poor, while simultaneously being highly targeted. Along the west coast of California many of the mesophotic fisheries are actively managed, with enforcement agencies frequently revising depth restrictions and species-specific take limits. Active management requires monitoring data from these depths and a combination of fisheries-dependent and independent data has been shown to provide a robust measure of stock size (Dennis et al. 2015; Howard et al. 2023). Baited remote underwater video (BRUV) systems have opened new opportunities for better understanding mesophotic reefs in temperate and tropical ecosystems by providing a fisheries-independent, non-destructive sampling technique that can operate at depths beyond recreational or even most scientific SCUBA diving limits (Brown et al. 2022; Andradi-Brown et al. 2016; Brokovich et al. 2008).

Mesophotic reefs represent an ecologically important habitat that contains distinct invertebrate and fish communities (Williams et al. 2019; Honeyman et al. 2023; Bell et al. 2024). Below 30m depth, solar irradiance begins to attenuate, limiting photosynthesis and algal growth that characterizes shallower reefs. At these depths, structure forming sessile invertebrate, many themselves of conservation concern, such as gorgonians, anemones, calcareous algae, and bryozoans dominate the benthos (Ponti et al. 2018). In the Northern hemisphere, the mesophotic temperate reef fish community is dominated by Rockfishes (genus *Sebastes*), a diverse assemblage of ecologically and economically important fishes that includes >100 recognized species (Love et al. 2002; Love and Yoklavich 2006; Love 2011). The mesophotic depth zone offers a relatively stable set of environmental conditions that can function as nursery habitat and a refuge space for species that were historically found along wider depth gradients (Baillon et al. 2012; Kahng et al. 2017; Giraldo-Ospina et al. 2020). These habitats also provide an important ontogenetic bridge between shallow nursery habitats and deeper adult habitats (Love 2011; Swadling et al. 2022). Finally, these deep habitats might be critical refuges from the effects of climate change, such as increases in sea water temperature (MacDonald et al. 2016; Pereira et al. 2018). Together these factors make mesophotic reefs important contributors to marine biodiversity and a necessary component of an effective MPA network.

One of the expected benefits of MPAs is the buildup of targeted species within its boundaries and consequently, an MPA’s performance is often measured by its ability to enhance the biomass and abundance of fisheries targets (Caselle et al. 2015; Lenihan et al. 2022). Recreational and commercial fisheries often have a number of targeted species that utilize a wide breadth of habitats across the seascape (Region 2003; Scholz et al. 2006).

Effective MPA networks should protect habitats ranging from sand, sand-rock ecotone, rocky reefs, and pelagic zones, while incorporating these habitats across an extensive depth range (Young and Carr 2015). Despite the fact that MPAs are an ecosystem conservation tool, it is unlikely any MPA can achieve a ‘one-size fits all’ design, instead, any single MPA will likely be more effective at protecting some species compared to others. Factors such as species movement, the availability of preferred habitat, oceanographic conditions, predator and prey abundances, and recruitment dynamics will all influence an MPA’s ability to enhance particular targeted species (Moffitt et al. 2011).

With the purpose of better understanding the effects of MPAs on mesophotic rocky reef fish communities, we implemented the use of BRUV camera systems at two geographically distinct MPAs (and nearby reference areas) within California’s Northern Channel Islands (Figure 1): Carrington Point State Marine Reserve (SMR) at Santa Rosa Island and Anacapa Island SMR/State Marine Conservation Area (SMCA). Straddled across a biogeographic transition zone, where cool southern-flowing waters of the California Current meet warmer northern-flowing waters from the Southern California Counter Current (Horn and Allen 1978), these two MPAs are characteristically distinct, occupying opposite ends of a strong environmental gradient that exists across the Northern Channel Islands. The primary variation along this gradient is in sea surface temperature (SST), accompanied by other related factors including productivity, frequency of disturbance/wave exposure, algal species persistence, and level of human interaction (Harms and Winant 1998; Hamilton et al. 2010; Caselle et al. 2015).

**Figure 1.**
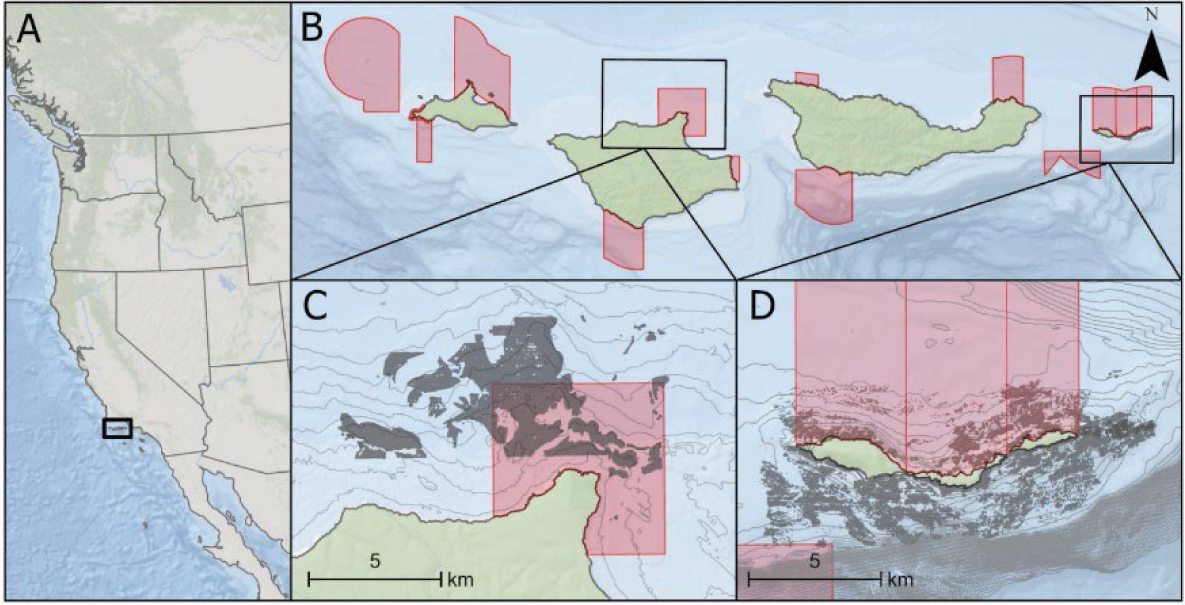
Maps showing the study region and the two focal MPAs in the Santa Barbara Channel, Northern Channel Islands (NCI). A) shows the location of the NCI along the West Coast of North America. B) shows the NCI with red polygons delineating the NCI MPAs within the state-wide California Marine Protected Area Network. Panel C) shows Carrington Pt. MPA and associated reference area while D) shows Anacapa MPA complex and associated reference area. Both panels C) and D) have dark grey shading representing mapped hard bottom and 10m bathymetry lines in light grey. Locations of BRUV surveys by year are shown in Figure S2.

We utilized stereo BRUV surveys because they are well suited for capturing fish community data across this under-sampled depth range and are useful for a variety of habitat types (Cappo et al. 2004; Heyns-Veale et al. 2016; Whitmarsh et al. 2017). Additional advantages of assessing fish communities using BRUVs include its non-destructive nature as a sampling technique (essential for MPA monitoring), the ability to collect highly accurate fish measurements for biomass calculations, the creation of a video record that can be re-analyzed to extract novel information or correct for inter-observer biases, the ability to observe species targeted by fishing and non-targeted species, and the opportunity to collect visual data on species habitat associations (Cappo et al. 2006; Langlois et al. 2010; Bennett et al. 2016). BRUV systems have become a widely implemented technique in many parts of the world as a non-destructive, fishery independent data collection tool for monitoring marine protected areas (Rees et al. 2004; Langlois et al. 2006; Kelaher et al. 2014) but are still in the early stages of use in California.

Using BRUV survey data from Carrington Pt. SMR and Anacapa SMCA/SMR and nearby reference areas we measured the importance of habitat type, depth, and protection status on the abundance and biomass of targeted fish species. These two focal MPAs are nested into larger networks; first, a network within the Channel Islands (Figure 1B), which itself is nested into a large California statewide network of protected areas. Previous studies have shown individual variation in MPA performance across the network for shallow reefs and kelp forests (Caselle et al. 2015; Ziegler et al. 2023). Here we use two MPAs to demonstrate the potential for individual MPA variation in performance in presumably more stable, deeper water habitats.

## Methods

### Sampling Design

Our BRUV design was modified from existing designs used extensively to monitor a variety of marine habitats in Australia (Willis and Babcock 2000; Goetze et al. 2021; Harvey et al. 2021). We deployed BRUVs using a stratified random approach, using geological habitat maps and bathymetry to target rocky reef habitat within our sampling depths (30-100m). Because the Carrington Pt. SMR has very little habitat >50m deep, survey depths at this site were generally limited to this maximum depth, while surveys at Anacapa extended to 100m depth. A map showing each MPA/reference zone polygon can be found in Figure S1.

Sampling effort included both MPA and reference sites (where fishing is allowed) on any single day to control for environmental variability between sampling days. Sampling took place from August to October for four consecutive years (2019-2022). In 2020, we also sampled in June at Anacapa Island as a response to the shutdown of commercial passenger fishing vessels (CPFV) at that location due to COVID-19. We tested for seasonal or COVID related affects and finding none, this data was included in all analyses unless otherwise noted. A map of all sampling locations, by year is available in Figure S2. A more detailed description of BRUV sampling methods and tools can be found in supplemental materials Methods S1.

### Video Processing

We analyzed video files using observation logging and 3D measurement software SeaGIS EventMeasure (www.seagis.com.au). We identified all observed fishes to the lowest taxonomic group possible. In order to quantify the relative abundance of fishes we recorded MaxN, the maximum number of individuals of a species present in a single video frame (Willis and Babcock 2000) using the first 30 minutes of each BRUV survey (Harasti et al. 2015), for every species observed. Without the ability to discern individuals or distinguish multiple sightings of the same fish, MaxN is a conservative estimate of relative abundance that has become the standard metric for BRUV surveys (Langlois et al. 2020).

We also measured the total length for every fish observed in the MaxN video frame. In some instances, we were not able to accurately measure all individuals in a particular MaxN frame due to the camera angle or obstructions blocking one of the cameras. In this case, we measured as many individuals as possible from the MaxN frame. We estimated the biomass of individual fishes using an allometric length-weight conversion: W = aTLᵇ, where parameters a and b are species specific constants, TL is total length in cm, and W is weight in grams. We obtained length-weight fitting parameters from the literature and FishBase (Froese and Pauly 2000). Because not all fish in the MaxN frame can be measured (some individuals are not within view of both cameras) we used the mean of all individual weights in the MaxN frame, multiplied by the MaxN for each individual species to estimate biomass. We classified species as targeted by fishing or non-targeted by fishing using information from the CA Department of Fish and Wildlife as well as local knowledge of the authors.

In order to understand the influence of habitat on MPA effectiveness, and because we cannot control the precise view of the forward facing BRUV cameras, we classified habitat based on substrate characteristics within the field of view of the cameras. The four habitat classifications were “Hard”, “Mixed-Hard”, “Mixed-Soft”, and “Soft” (Categories defined in Table S1).

To standardize the quality of videos used in analysis, we quantified the “usability” of each video. All videos were assigned a “usability” score that considered the amount of visible substrate and video length (Table S2). Any videos where the BRUV unit did not remain upright for the entire 30 min (i.e., scores of 3 or 4) were removed from analyses.

### Focal fish species

In addition to groupings of all targeted species and all non-targeted species, we also analyzed patterns of biomass and MaxN for specific focal species. Ocean whitefish, the most abundant species observed in our study, are a medium size (39-56 cm TL at sexual maturity) tilefish (*Malacanthidae*) common in Southern California and Baja, Mexico. Typically residing at the periphery of rocky reefs, ocean whitefish primarily feed on small invertebrates often in nearby sandy flats (Love 2011). This species is a staple for the recreational fishery in the heart of its geographic range and is likely to benefit from fishing protection.

California sheephead, which appeared on nearly half of our surveys, are the largest and only targeted wrasse (*Labridae*) species that inhabits the study area. California sheephead are not only highly targeted by recreational anglers, but also by a smaller live-fish commercial fishery (Love 2011). As a protogynous hermaphrodite (female at birth with the potential to transition to male later in life), fishing pressure has a more complex effect on the sex/size structure and the reproductive potential of a local population (Hamilton et al. 2007).

With 21 unique species of rockfish observed in this study, we more closely examined a sub-group of rockfish that we considered “targeted” and are more likely to respond to protection from fishing. This grouping excludes small-bodied rockfish that are infrequently caught and almost never retained by anglers such as halfbanded (*S. semicinctus*) and calico rockfish (*S. dallii*). Three targeted species: vermilion (*S. miniatus*), copper (*S. caurinus*), and blue rockfish (*S. mystinus*) made up over 70% of the targeted rockfish observed on our surveys. Vermilion and copper rockfish are long-lived (50-60 years) high value species that have a significant recreational and commercial fishery, increasing their potential to benefit from protection. Blue rockfish are a schooling, largely planktivorous species, that is of moderate value to recreational and commercial fishing (Love 2011).

### Abundance and Biomass Modeling

Generalized linear models (GLM) were used to understand how differences in habitat type, depth, and MPA status influenced fish abundance and biomass. To account for species-specific habitat preference and other potential traits (e.g. movement patterns) we ran models on targeted rockfish, ocean whitefish, and California sheephead separately. We also ran separate models for each island because the depth ranges differed (Carrington Pt. SMR ranged from 30m - 60m depth; Anacapa SMCA/SMR ranged from 30m – 90m depth). Predictor variables were MPA status, depth, and habitat type, as well as the interaction between MPA status and habitat type to test if the effect of MPA status is similar across different habitat types.

All statistical models and model diagnostics were run using R version 4.3.0. Data from each species and metric (biomass and MaxN) combination was zero inflated, and the fitdistrplus package and descdist() function were used to determine the best fit distribution. Each data combination was fit to a negative binominal distribution and the models were run using the glm.nb function in the lme4 package. Randomized quantile residuals were calculated and tested for overdispersion using the DHARMa package and all dispersions statistics were <1. The ‘visreg’ package was used to visualize how each model parameter influenced a particular metric and represents each model estimate when all other predictors are held constant. We standardized the visualizations by setting MPA status to ‘MPA’, habitat type to ‘Hard’, and depth as the median depth from a particular island. Each regression plot shows a model prediction, 95% confidence interval, and the partial residuals of the model. We calculated the precent deviance explained for each model to understand how well our predictors explained the variation in the data.

### Population Size Structure

To understand how protection from fishing alters the demographics of targeted fish populations, we plotted the size density histograms of the three most common targeted rockfishes (copper, vermillion, and blue rockfish), ocean whitefish, and California sheephead.

## Results

We completed 280 and 341 BRUV surveys at Carrington Pt. and Anacapa Island respectively for a total of 621 surveys. Of those 621 surveys, 493 were classified as ‘usable’ (79% usable) and were included in analyses (Table S3) In general, two conditions resulted in unusable video; these were very strong currents and/or highly rugose habitat causing the BRUV system to not land upright, or tip over at some point during the video survey.

### Habitat Description

We found differences in amount and type of rock at the two islands. When rock was present at Anacapa, the habitat was characterized by a patchy reef system with most video classified as ‘mixed soft’ or ‘mixed hard’ (Fig. 2, Table S4). In addition, the Anacapa MPA contained more ‘soft’ habitat classifications than the other island/MPA groupings. Carrington Pt. was characterized by more consistent hard bottom, with most videos classified as ‘hard’ or ‘mixed hard’ (Figure 2, Table S4).

**Figure 2.**
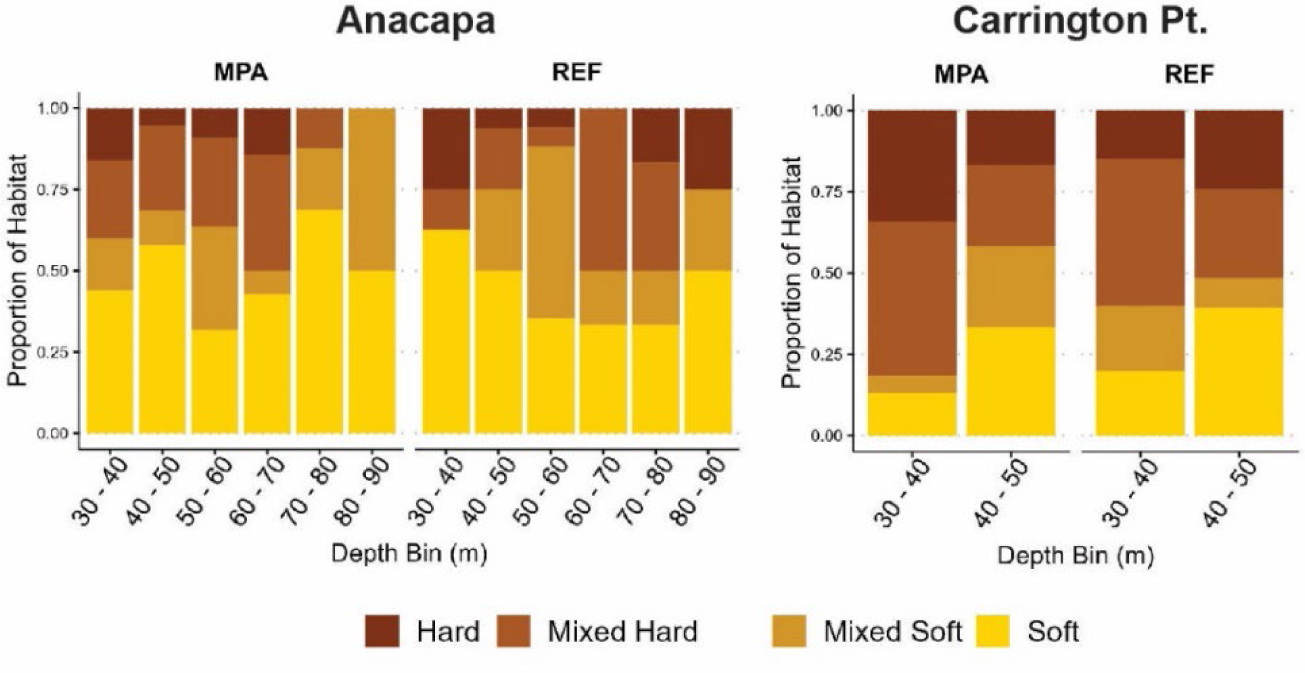
Stack bar plot showing the proportion of observed habitat by depth bin from Anacapa and Carrington Point. Habitat was classified from BRUV surveys and was categorized based on visible habitat in which a survey was conducted.

Similar to the video observations, benthic habitat maps showed a disparity between the amount of rocky substrate present in the Anacapa MPA and reference area, with 3-5 times more eligible grid cells in the reference area. Although the reference area was twice as large as the MPA, there still appears to be proportionally more abundant hard bottom habitat in the reference area. This is particularly true for the deepest zone (70-100m), which had 5 times as many eligible grid cells (MPA = 33 vs Reference = 167, Table S5) in the reference area. Carrington Pt. had a relatively similar number of eligible grid cells within the 30m - 50m depth bin, with the reference area only being slightly larger (Table S5). Overall, the benthic habitat maps used to identify eligible grid cells generally agreed with the observed habitat data from BRUV surveys, with Carrington Pt containing more consistent hard bottom and Anacapa containing more sand-rock ecotone habitat.

### Species Observed

We observed a total of 65 fish species on the BRUV surveys across four years and both locations. Total summed MaxN and frequency of occurrence for each species are shown in Table S6. A single species, ocean whitefish, accounted for 32% of the summed MaxN values. These fish had the highest abundance across our study and were present on 84% of all surveys. Two species, copper rockfish and California sheephead had both high MaxN values and high frequency of occurrence at both islands. Copper rockfish occurred on 48% of surveys while California sheephead occurred on 49% of surveys (Table S6). Four schooling species: jack mackerel (*Trachurus symmetricus*), blacksmith (*Chromis punctipinnis*), halfbanded rockfish, and blue rockfish were abundant when observed but occurred less frequently (4-27% of all surveys).

The BRUVs were able to observe species rarely seen in other surveys in California (e.g., SCUBA, ROV, submersible). These included species of concern including giant sea bass (*Stereolepis gigas*) (50 individual observations), bocaccio (*Sebastes paucispinis*) (79 individuals) and bat rays (*Myliobatis californica*) (11 individuals) as well as various species of the more cryptic flatfishes (including California halibut (*Paralichthys californicus)* and Pacific sanddabs (*Citharichthys sordidus*)). Although many species were documented by only a single individual, this type of presence-only data can be useful for documenting climate-induced range shifts, invasions, and habitat associations.

### Abundance and Biomass

#### Targeted Rockfish

At both islands, targeted rockfishes showed similar patterns with habitat type and depth being significant predictors of both biomass and abundance. Generally, both biomass and abundance increased as the availability of hard substrate increased, with ‘hard’ vs. ‘mixed soft’ and ‘soft’ being highly significant (all eight comparisons across island, metric, hard vs. mixed hard, and hard vs. soft had a p-value <0.001; Figure 3 & Table S7). At Anacapa, depth was a significant predictor for targeted rockfish abundance and biomass, with both metrics increasing with depth (MaxN p-value = <0.001; biomass p-value = <0.001; Figure 3 & Table S7). The patterns at Carrington Pt. were less consistent, with depth only being a significant predictor of biomass (p-value = 0.02, Figure 3 & Table S7), most likely related to the limited depth range at this location compared to Anacapa. Interestingly, the interaction between soft habitat and MPA status was almost significant for targeted rockfish biomass at Anacapa. Here, soft habitat was the only habitat type predicted to have greater targeted rockfish biomass inside the MPA compared to the reference area (p-value = 0.06, estimate = −1.43; Table S7 & Figure S3A). This interaction term indicates that the model estimate for soft bottom habitats in the MPA was greater compared to reference areas, while in other habitats (e.g., ‘Hard’, ‘Mixed Hard’, and ‘Mixed-soft’) the reference area estimate was larger. The only significant MPA effect, beyond the influence of habitat and depth, was for targeted rockfish biomass inside the Carrington Pt. MPA (p-value = 0.03; Figure 3 & Table S7).

**Figure 3.**
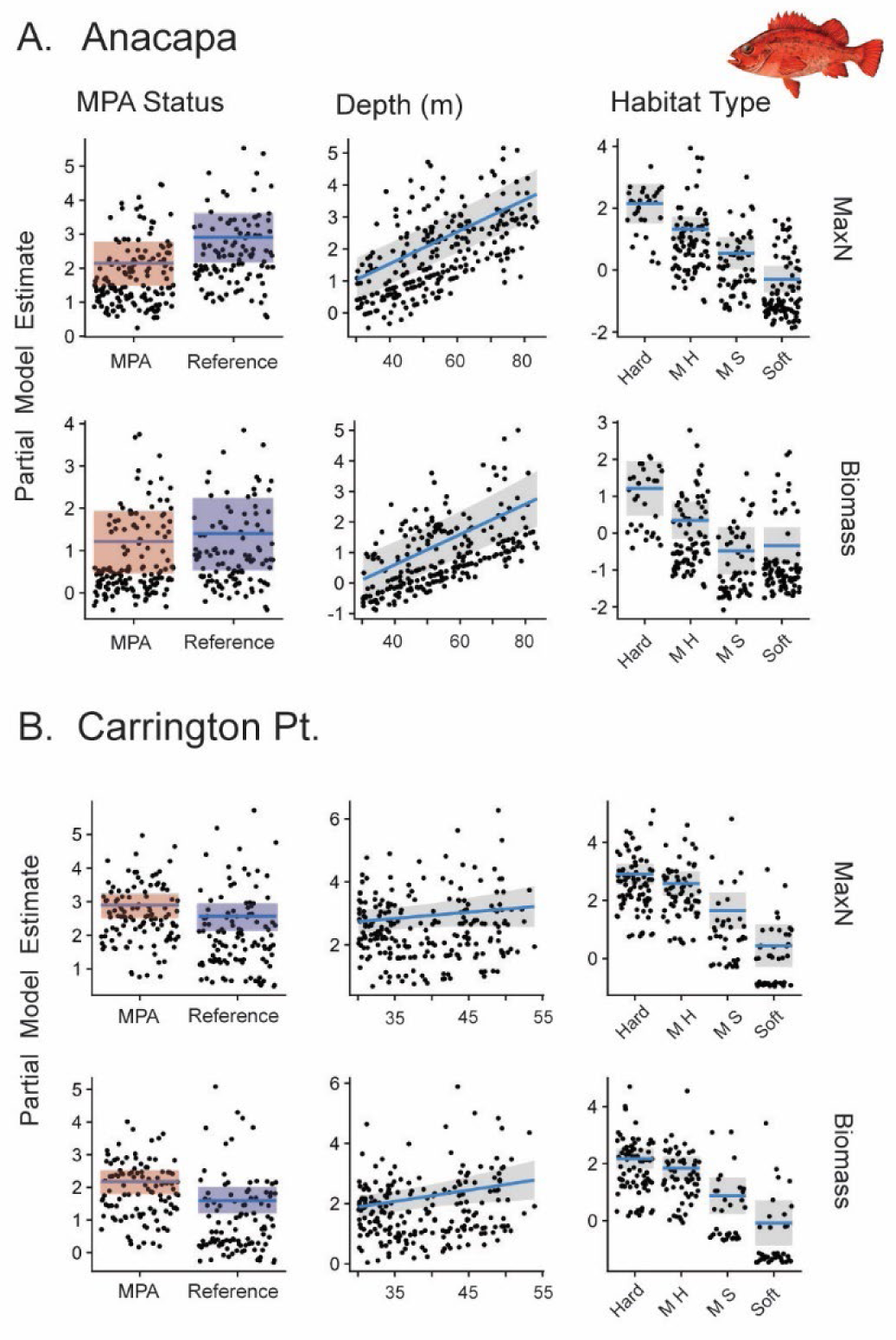
Partial model predictions from a negative binomial generalized linear model used to test the effect of MPA Status, depth, and habitat on the MaxN and biomass of targeted rockfishes for Anacapa (A) and Carrington Point (B). The blue line and shaded region represent the model predictions and 95% CI for a particular predictor variable when all other variables are held constant. The points represent the partial residuals of the model. The habitat codes ‘M H’ and ‘M S’ represent mixed hard and mixed soft habitats respectively.

#### Ocean Whitefish

Depth was a strong predictor of ocean whitefish biomass and abundance with higher abundance and biomass found in shallower waters (p-value = < 0.001 for all comparisons of MaxN and biomass at both islands; Figure 4 & Table S8). The only significant habitat predictor was for soft habitats at Carrington Pt., with both abundance and biomass significantly lower on sandy, soft bottom habitats compared to hard substrate (p-value = 0.05; Figure 4 & Table S8). We did not find a significant interaction term between MPA status and habitat type for either ocean whitefish metric (Figure S4 & Table S8). Anacapa showed a significant MPA effect, with the abundance and biomass of ocean whitefish greater inside the MPA (MaxN p-value = 0.02; biomass p-value = 0.01; Figure 4 & Table S8).

**Figure 4.**
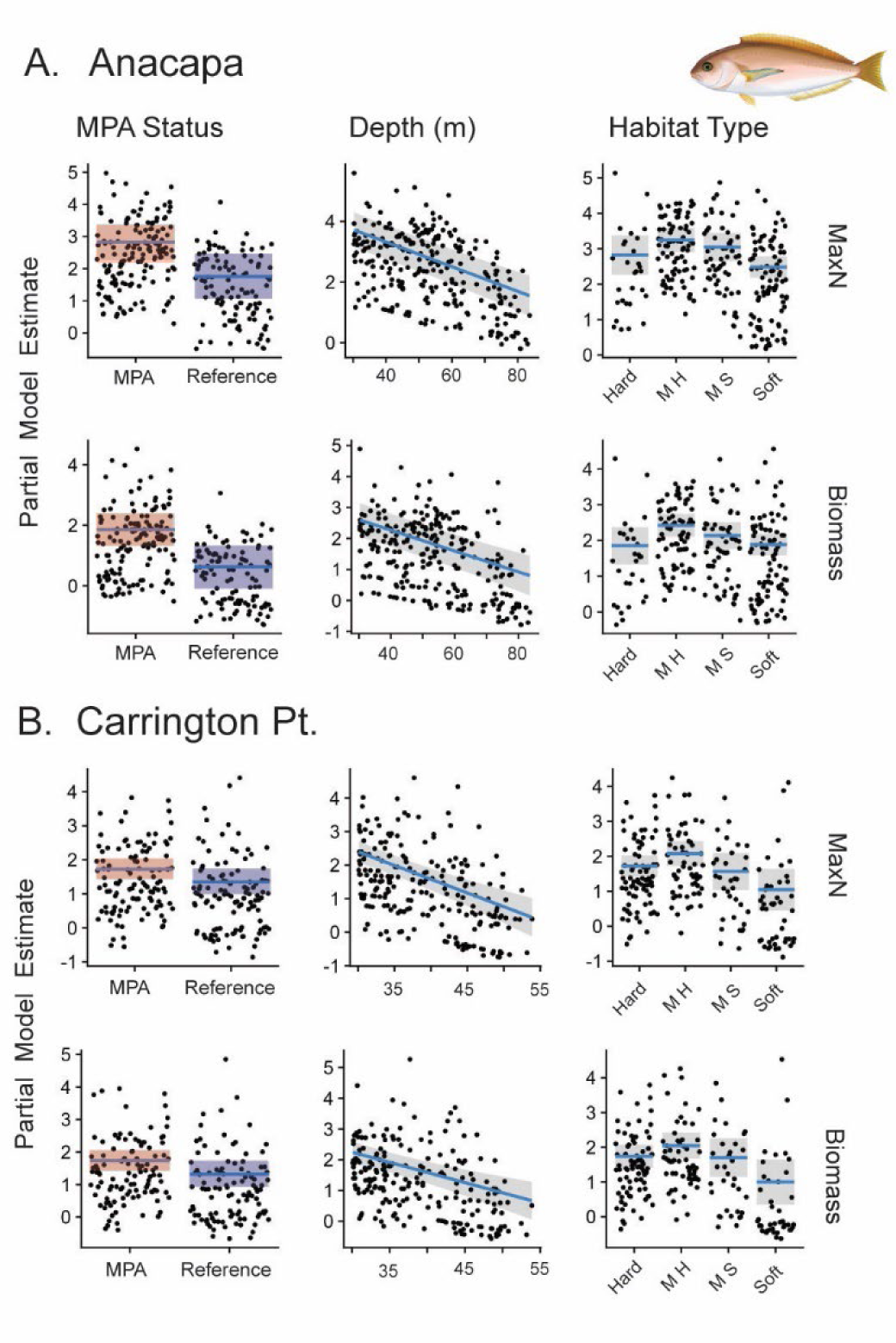
Partial model predictions from the negative binomial generalized linear model used to test the effect of MPA Status, depth, and habitat on the MaxN and biomass of ocean whitefish for Anacapa (A) and Carrington Point (B). The blue line and shaded region represent the model predictions and 95% CI for a particular predictor variable when all other variables are held constant. The points represent the partial residuals of the model. The habitat codes ‘M H’ and ‘M S’ represent mixed hard and mixed soft habitats respectively.

#### California Sheephead

Similar to ocean whitefish, depth was found to be a strong predictor of California sheephead biomass and abundance, with higher levels found in the shallower waters (Carrington Pt. and Anacapa MaxN p-value <0.001; Carrington Pt. biomass p-value = <0.01; Anacapa biomass p-value = <0.001; Figure 5 & Table S9). California sheephead were significantly less abundant on two habitat types at Anacapa, ‘mixed soft’ and ‘soft’, compared to ‘hard’ substrate (‘mixed soft’ p-value = 0.03; ‘soft’ p-value = <0.01; Figure 5 & Table S9), and had significantly lower biomass on ‘soft’ vs ‘hard’ habitat at Anacapa (‘soft’ p-value = 0.05; Figure 5 & Table S9). At Carrington Pt., habitat alone was not found to be a significant predictor of sheephead abundance or biomass. We did find a significant interaction term for sheephead biomass and abundance between ‘soft’ habitat and MPA status at Carrington Pt., however these results had very few observations of sheephead on soft habitat across protection zones, limiting inference from this result. At Anacapa, we found a significant negative MPA effect for the abundance and biomass of sheephead, with greater values in the reference area (MaxN p-value = 0.01, estimate = 1.01; biomass p-value = 0.03, estimate = 1.09; Figure 5 & Table S9).

**Figure 5.**
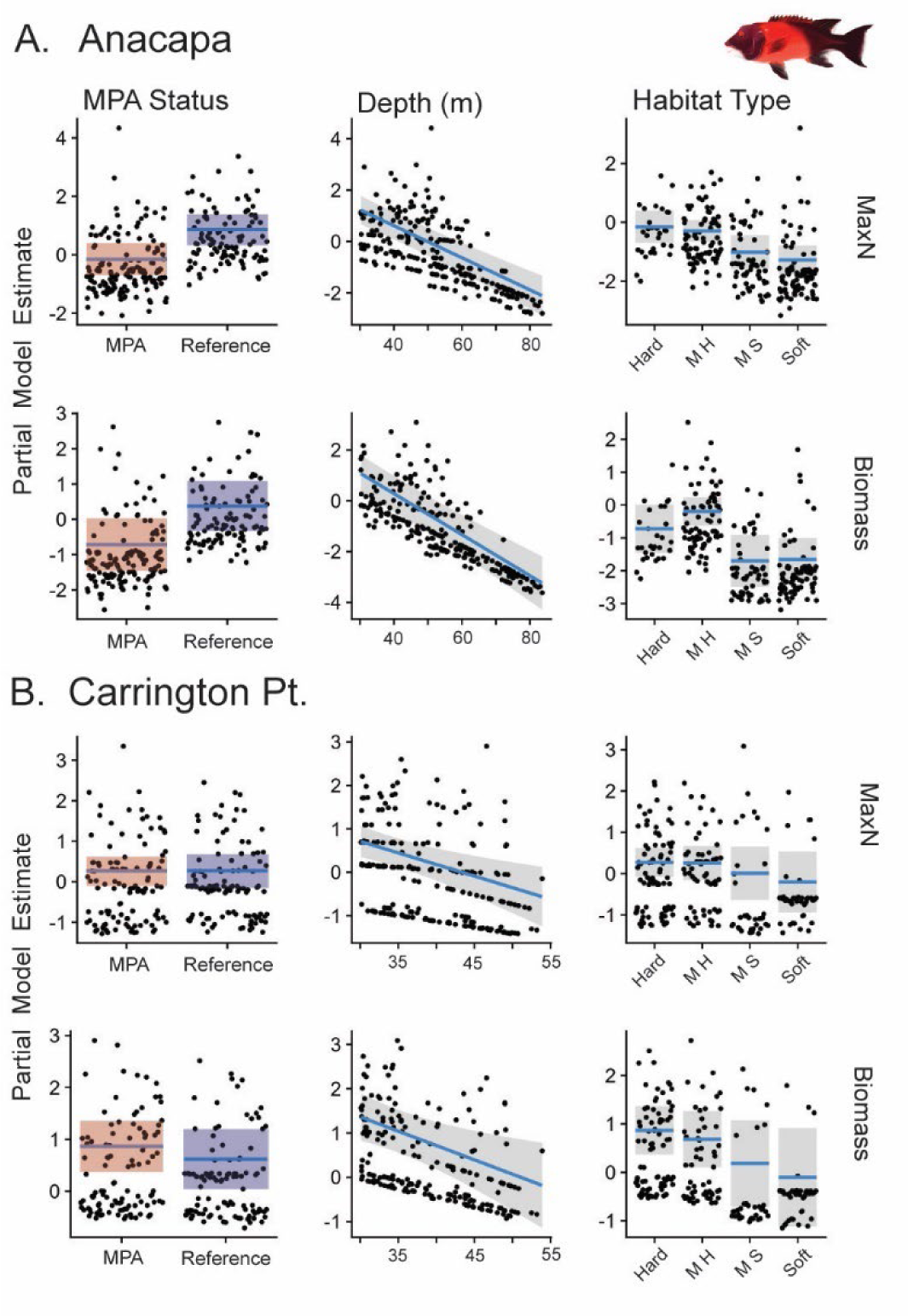
Partial model predictions from the negative binomial generalized linear model used to test the effect of MPA Status, depth, and habitat on the MaxN and biomass of sheephead for Anacapa (A) and Carrington Point (B). The blue line and shaded region represent the model predictions and 95% CI for a particular predictor variable when all other variables are held constant. The points represent the partial residuals of the model. The habitat codes ‘M H’ and ‘M S’ represent mixed hard and mixed soft habitats respectively.

#### Deviance Explained

Across all species and both metrics, the model framework performed slightly better at Anacapa compared to Carrington Pt., with the deviance explained ranging from 0.46 – 0.22 at Anacapa and 0.22 – 0.17 at Carrington Pt. (Tables S7 – S9). This is likely related to the larger available depth range at Anacapa and that depth was a significant predictor of abundance and biomass for all three species in this study. The highest deviance explained was for California sheephead at Anacapa (DE = 0.46 for MaxN; DE = 0.45 for Biomass; Table S9) and is most likely related to the smaller range in MaxN and biomass values associated with the California sheephead data compared to the larger range associated with targeted rockfish and ocean whitefish.

### Focal Species Size Structure

We plotted the density of individual fish total lengths in MPAs and reference areas for the five most common targeted species in our study; copper rockfish (n = 506), vermilion rockfish (n = 459), blue rockfish (n = 1760), ocean whitefish (n = 5767), and California sheephead (n = 603) (Figure 6). Copper rockfish were generally larger inside the MPAs at both Anacapa and Carrington Pt., with the MPA containing more individuals >400mm TL (Figure 6A). Vermilion rockfish had a bimodal size structure with multiple size classes present at Carrington Pt. and Anacapa (Figure 6B). Inside the Anacapa MPA, the population size structure is dominated by smaller sized individuals (<20cm), with proportionally fewer adult fishes. While the population size structure patterns for vermillion rockfish at Carrington Pt. appear to be more related to protection from fishing, with proportionally larger individuals inside the MPA. The smaller-bodied schooling blue rockfish had similar population size structure across protection zones at Carrington Pt. and Anacapa (Figure 6C). Ocean whitefish and California Sheephead showed similar patterns at Anacapa and Carrington, with a greater proportion of larger individuals inside the MPA at each location (Figure 6D & E).

**Figure 6.**
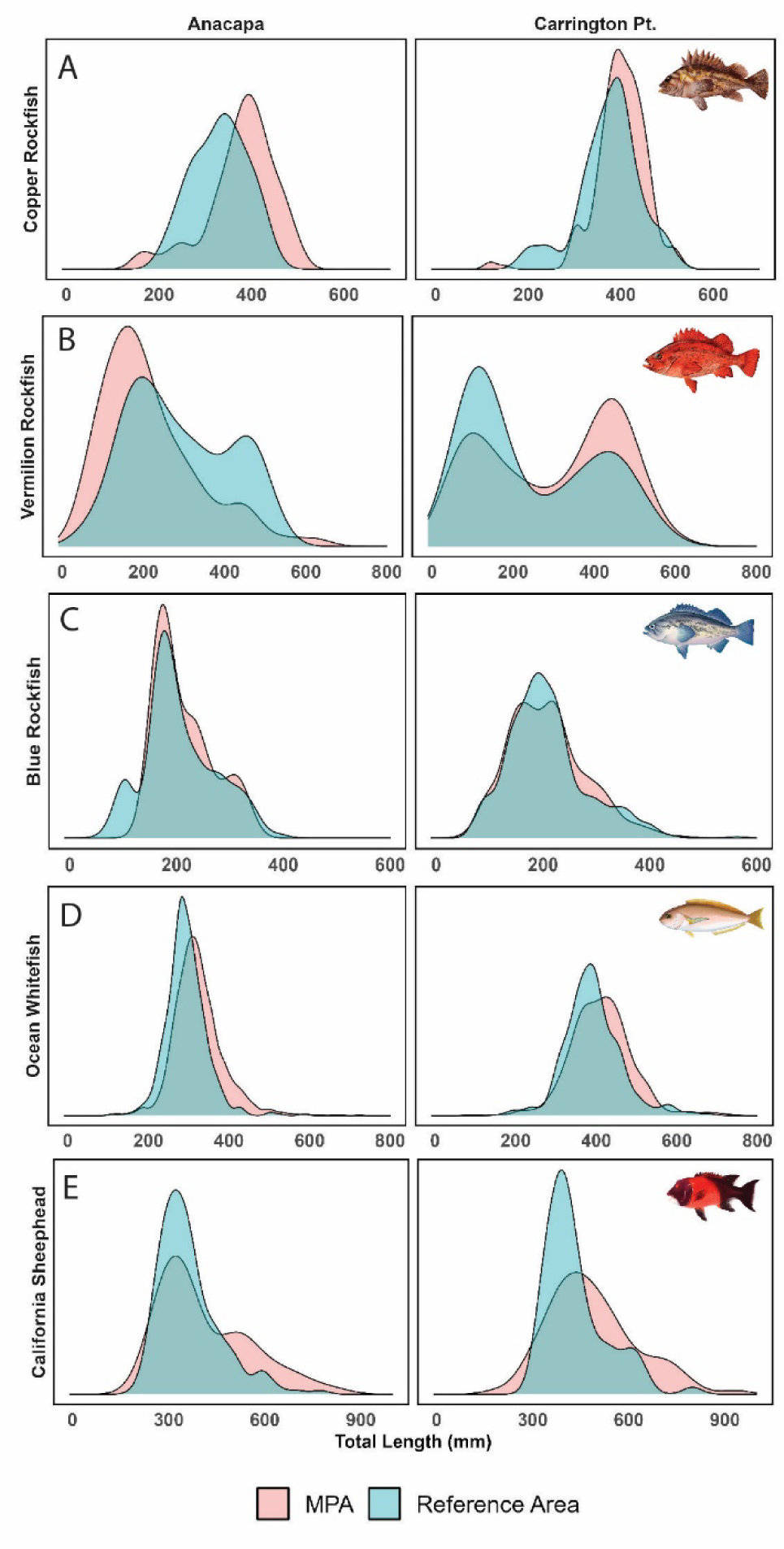
Species-specific density histograms for the five targeted fish species; Copper Rockfish (A), Vermillion rockfish (B), Blue rockfish (C), ocean whitefish (D), and California sheephead (E) from Anacapa and Carrington Pt. The red density histograms represent data from within the MPA, while blue density histograms represent the data form reference areas open to fishing.

## Discussion

In this study we show that environmental drivers, such as depth and habitat availability, need to be considered when assessing the ability of MPAs to enhance the biomass and abundance of targeted species. MPA effectiveness, in both tropical and temperate marine ecosystems, has been linked to benthic features such as the amount of consolidated hard bottom, reef complexity, and habitat continuity (Dames et al. 2020). Individual species have been shown to respond differently to such features, hence species-habitat associations must be considered during MPA implementation, when defining MPA goals, and when assessing performance.

Studies have shown that a lack of preferred habitat inside the protection zone can limit the ability of MPAs to enhance species and functional diversity (Navarro-Martínez et al. 2023). MPAs are designed as ecosystem management tools, with broad goals of protecting biodiversity, yet individual species can have widely varying habitat requirements. Many MPA design processes, especially for MPA networks, try to encompass habitat replication and representation (Roberts et al. 2003; Halpern et al. 2010; Saarman et al. 2013), but in practice, it is impossible to design optimally for every species. Thus, accounting for habitat in post implementation monitoring and choice of reference areas, can help to set realistic expectations for MPA performance.

The Carrington Pt. MPA and adjacent reference area had similar amounts of continuous hard substrate, which indicated that the greater biomass of targeted rockfishes in the MPA was likely related to protection more than habitat. Rockfish (family *Sebastes*) are generally characterized as a demersal, rock associated and heavily fished family, and it appears that the Carrington Pt. MPA is well designed for their protection. Inside the Anacapa MPA, there was a notable lack of continuous hard substrate, particularly at depth, that appears to limit the performance of the MPA at that location for this rock-associated fish group and makes comparisons with the paired reference area challenging. However, the Anacapa MPA is likely more effective at enhancing the ocean whitefish population, a more mobile species that is associated with the sand-rock ecotone (Bellquist et al. 2008). Habitat maps and *in situ* observations from Anacapa MPA and reference area showed that these locations were dominated by a mix of soft, sandy habitat and isolated patch reefs. Ocean whitefish abundance and biomass was also strongly associated with depth, but unlike targeted rockfish, ocean whitefish are more abundant in shallower areas, with the majority of observations occurring shallower than 60m.

The examples above, comparing very different fish species groups, demonstrate how scientific guidelines and a network approach to MPA design can help meet ecosystem goals and protect populations of different species across a variety of habitats. The creation of networks, with numerous reserves spread across various habitats, has become the standard by which to structure marine no-take zones, both at the local and global scale. The California Marine Life Protection Act (MLPA) established the network of MPAs throughout California with the goals of protecting ecosystem biodiversity and protecting and rebuilding populations of key species. This process relied on MPA design guidelines from a scientific advisory committee and chief among those guidelines was ‘habitat representation’ throughout the network (Saarman et al. 2013). At the network level, this was achieved by including a variety of major habitat types such as; rocky intertidal zones, estuaries, and shallow and deep subtidal habitat. Here we show that a more nuanced treatment of habitat within an MPA can provide critical information on individual MPA performance for key species (Young and Carr 2015).

By incorporating habitat availability in and out of the MPAs and studying deeper zones than previously explored, we found several results that differed from prior studies and/or our expectations. Contrary to our predictions and prior studies of shallower reefs in the area (Caselle et al. 2015; Honeyman et al. 2023), California sheephead abundance and biomass at these deeper depths was significantly greater in the Anacapa reference area which is open to fishing. Active tracking data has shown that California sheephead use a variety of habitats, but tend to favor rocky-reefs, spending 50% - 70% of their time on hard, consolidated substrate (Topping et al. 2005). The deep-water habitat within the Anacapa MPA is lacking this hard substrate, which is more abundant in the shallow kelp forests of the Anacapa MPA. This illustrates the importance of surveying the full range of depths for species of interest when evaluating MPA performance or advising fisheries management. A more comprehensive approach to MPA design and management can be achieved by filling gaps in our knowledge of how MPAs may differentially enhance fish communities at all depths and habitat types.

To measure the ecological effects of a marine protected areas, especially without data for that site prior to MPA implementation, researchers often use a nearby fished area (reference site) as a control by which hypotheses about the cessation of fishing can be tested. Selection of these ‘control’ sites must weigh numerous factors such as; habitat type, depth range, species composition, and seasonal oceanographic characteristics, when deciding on location and scale. Often the easiest way to broadly control for various environmental factors is by placing the reference area in close proximity to the MPA. However, unlike strictly controlled lab experiments, MPAs are expected to affect control areas by design. Adjacent placement of reference areas, while often meeting goals of controlling for environmental conditions and perhaps even habitat, creates a scientific design challenge, with the likelihood that reference sites will be affected by the presence of a nearby MPA through the redistribution of fishing effort, and potential “spillover” of fish coming from the MPA (Caselle et al. 2015; Di Lorenzo et al. 2016).

Anacapa Island presents an interesting case study in reference site selection. Previous studies of very deep water habitats using Remotely Operated Vehicles excluded the Anacapa MPA from analyses, citing a lack of appropriate reference area (Karpov et al. 2012). This exclusion comes despite containing one of the oldest MPAs in the Northern Channel Islands network, with the Anacapa eastern SMR established in 1978. Here we chose the reference location as immediately offshore of, and deeper than, long-standing reference sites used for shallower rocky reef and kelp forest studies (Caselle et al. 2015; White et al. 2021). While shown to be appropriate for shallow rock reef, we found that in deeper waters, the reference area contained more deep rock than in the MPA, potentially limiting our ability to detect an MPA effect on rock-associated species (e.g., targeted rockfish, California sheephead). However, both reference area and MPA contained similar amounts of deep sand-rock ecotone habitat favored by ocean whitefish, the most numerous species of fish in our study. Hence tradeoffs exist where reference site selection must incorporate species-habitat associations, assess MPA goals, work to inform management decisions, and are beholden to the realities of field studies.

A consistent MPA effect was observed in the size structure of three targeted rockfish species, ocean whitefish and California sheephead, with the greater proportion of larger individuals inside the protection zone. This is not surprising as one of the earliest effects of a cessation of fishing is to allow larger individuals to grow and persist (Mumby et al. 2021; Bejarano et al. 2019; Taylor and McIlwain 2010). The notable exception was for vermillion rockfish inside the Anacapa MPA, which contained fewer large individuals than the reference area. Interestingly, the size structure for vermilion rockfish at both study sites reveal distinct cohorts at two size classes. This likely indicates periodic strong recruitment/survivorship for this species and provides an opportunity to track these cohorts over time. Repeated annual sampling such as done here allows the tracking of these types of population demographics through time to better understand year to year variability in recruitment dynamics for valuable fishery species. The ability to collect size structure data, with tools such as stereo BRUVs, can provide important additional insight into the effects of protection on these populations and is particularly important for the management of long-lived species like rockfishes.

The ability of MPAs to conserve and rebuild fish communities has been shown to improve with the protection of habitat across a wide range of depths (Curley et al. 2002; Goetze et al. 2021). Many species of fish utilize different depth zones and habitats across life stages, often with juvenile fish recruiting to relatively protected shallow nursery habitats and moving to deeper offshore habitats as they mature (Gibson et al. 2002; Aburto-Oropeza et al. 2009; Li et al. 2022). This ontogenetic movement is common with a number of the rockfish species observed in this study (Love et al. 1991) and should be considered in the context of MPA performance. The availability of suitable habitat along the ontogenetic depth profile likely influences the distribution of rockfish at varying life stages. Additionally, seasonal spawning migrations across depth gradients are observed globally and emphasize the significance of providing protection across this gradient (Luo et al. 2020; Lombardo et al. 2020; Thorburn et al. 2021). We observed a distinct break in suitable rockfish habitat within the Anacapa MPA complex, with very little deep habitat (>70m) present. Not only does this reduce the overall available habitat for adult rockfish, it likely reduces the potential for this MPA to provide a valuable ontogenetic bridge for species with high habitat affinity. An MPA that contains contiguous habitats within a species full depth range is better equipped to protect that species at all stages of life, enhancing the benefits experienced by protection from fishing.

The protection of connected depth gradients is also likely to provide resilience for fish populations faced with environmental stressors including seasonal changes in water temperature, short-term temperature anomalies (marine heat waves), and long-term changes resulting from global climate change. In an ever-warming global ocean, it has become necessary for fish communities to adapt to rising sea temperatures, often through range shifts in latitude or depth (Perry et al. 2005; Dahms and Killen 2023; Chaikin et al. 2022). Access to deep refuge habitat can improve the resilience of some fish communities and allow them to survive shallow water stressors (MacDonald et al. 2016; Pereira et al. 2018). Inversely, global climate change has been shown to reduce deep-water dissolved oxygen, creating upward pressure on fish communities escaping oxygen minimum zones (OMZ) (Ross et al. 2020). This has been observed around Southern California’s Channel Islands, with some rockfish communities shifting to shallower depths as a response to OMZ shoaling (Meyer-Gutbrod et al. 2021). Spatial protections such as MPAs can provide corridors of protected habitat across depth zones to allow for this type of adaptive migratory process. The combined effects of large-scale environmental stressors and species-specific life histories on habitat use across depth gradients have the potential to alter an MPAs effectiveness in the near and long-term.

We used the same model framework across two species and one species group to understand the importance of MPA status, habitat type, and depth in determining patterns of abundance and biomass. Across all models, the model performance indicated that these factors were important in explaining the observed patterns, however other aspects of the system could be incorporated to more accurately evaluate conservation efforts. Future studies will work to integrate factors such as distance to reserve edge, presence of other ephemeral habitats (e.g. ophiuroid or urchin barrens), and oceanographic parameters. BRUVs are particularly well suited to adaptive monitoring programs, in that they are deployable in multiple habitats, relatively inexpensive to operate and provide a video record that can be reassessed over time (Langlois et al. 2010; Whitmarsh et al. 2017).

The California MPA network will continue to work towards protecting marine biodiversity and populations of key fished species across the state. Our results demonstrate the realized benefits of a well-designed and well enforced protection network, with a nuanced look at how two geographically distinct MPAs might effectively protect different fisheries targets. We also highlight the importance of acknowledging and describing the limitations of particular MPAs when assessing an MPA network to align expectations with realistic outcomes. Continued monitoring efforts will provide valuable ecological knowledge as MPAs age and management efforts adapt to preserve fisheries in the face of climate uncertainty.

## Acknowledgements

This work would not be possible without funding support from the California Ocean Protection Council and California Seagrant (Grant Numbers R/MPA-43 and R/MPA-48). Special thanks to C. Honeyman, A. Parsons-field, J. Kolda, B. Brock, B. Behar, P. Campbell, W. Horstmeyer, N. Castaneda, S. Jaeger, K. McCaffrey, S. Urgoiti Crespo, G. Hansen, J. Harding, and M. O’Connell for assistance with fieldwork. Thank you to Y. Chen, Y. Sha, T. Hozdic, G. Simmons, J. Zounes, and J. Eisaguirre for video analysis. Thank you to S. Gaines, D. McCauley, R. Starr, J. Lindholm, and J. Todd for advice and insightful discussions. Thanks to A. Giraldo Ospina and C. Free for help with data analysis.

