## Appendix for "Understanding organism-habitat relationships and critically evaluating reference areas is key to marine protected area assessment"

Supplementary material

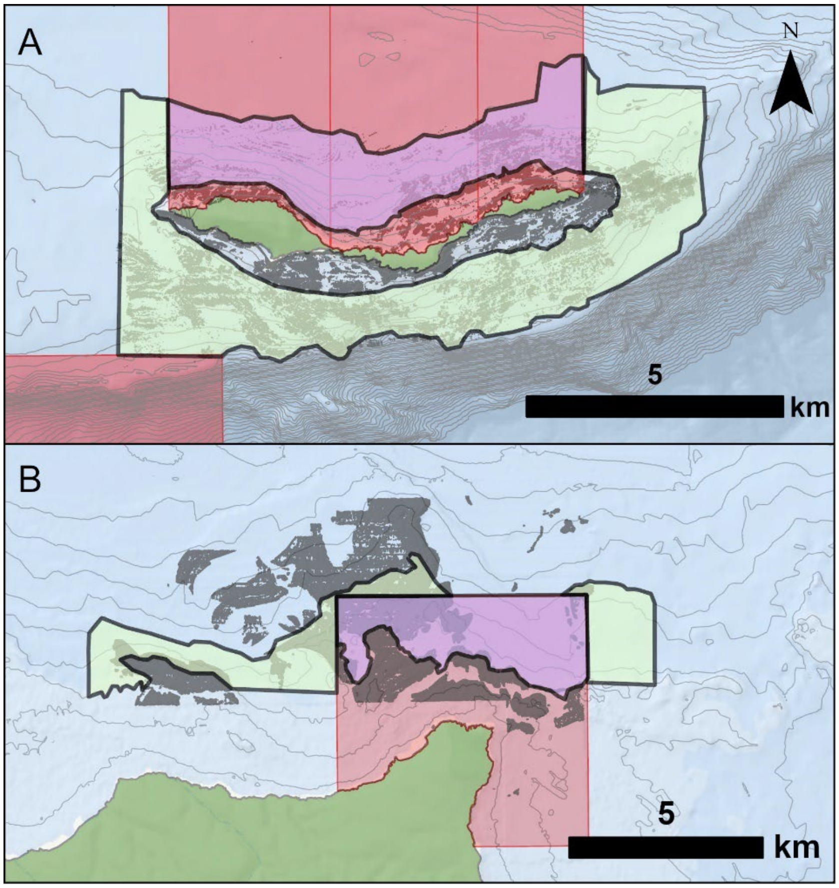


Figure S1. A) a map showing the Anacapa MPA complex and associated reference area, and B) a map showing the Carrington Pt. MPA and associated reference area. Each map shows the eligible study area within a particular MPA, represented by the purple polygon, and the survey area associated with reference area in the light green polygon. The dark grey shading represents mapped hard bottom used to identify survey locations and light grey lines representing 10m isobaths.

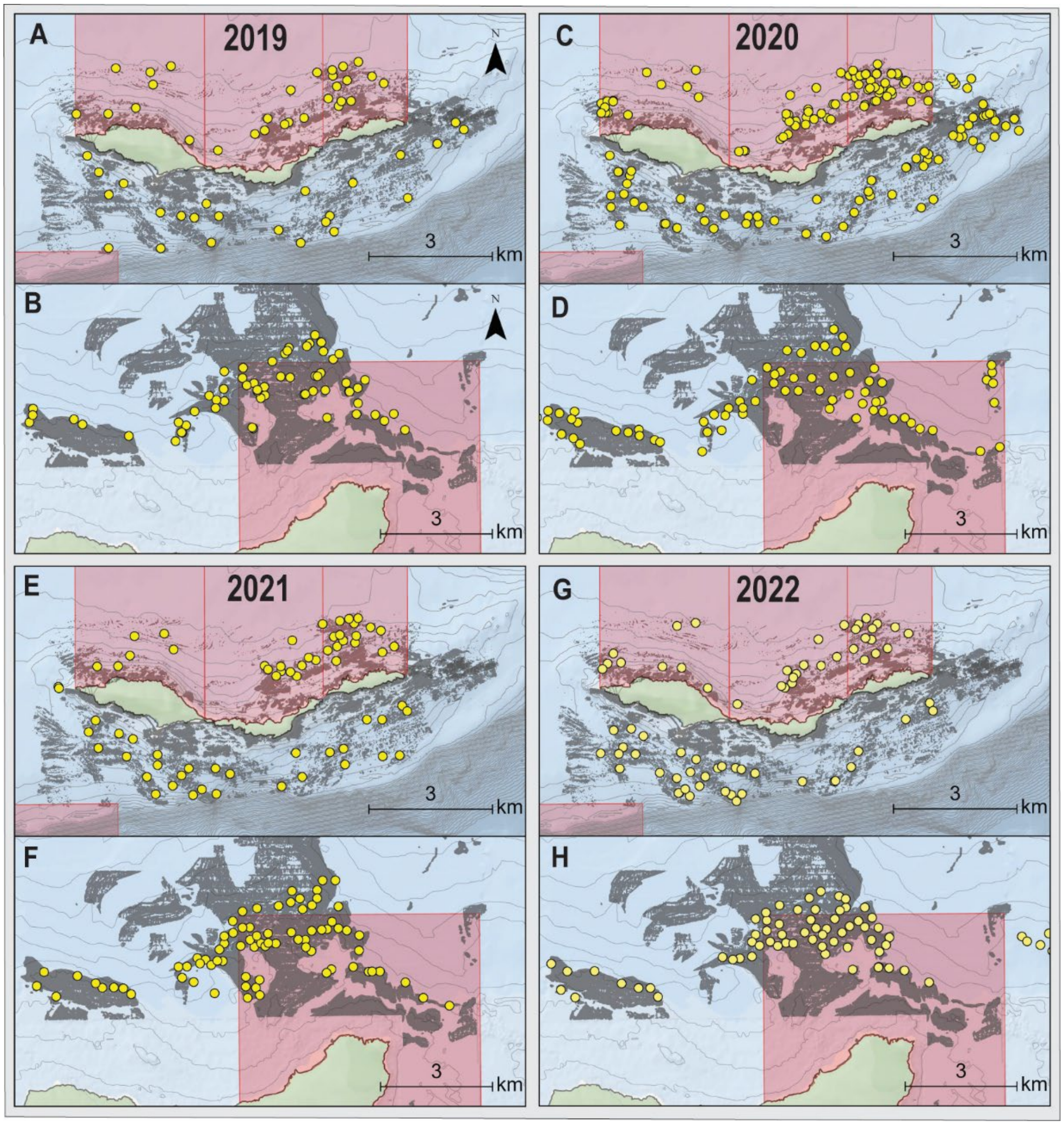


Figure S2. A set of maps showing the specific location of each survey in this study. The four panels each represent a particular year from 2019 to 2022. The upper map in each panel displays surveys at Anacapa and the lower map displays surveys at Carrington Pt. The shaded grey regions represent mapped hard bottom and the grey lines represent the 10m isobath.

Table S1. Definitions for habitat classification categories from BRUV survey videos.

| Habitat Type | Description |
|---|---|
| Hard | Substrate primarily consists of rock, boulders, or bedrock (<25% sand) |
| Mixed Hard | Substrate consists of rock and boulders (>25% sand) |
| Mixed Soft | Substrate consists of low relief bedrock/cobble and no boulders (>25% sand) |
| Soft | Substrate is entirely sand, no rock visible |

Table S2. Definitions for video usability scoring.

| Usability Score | Landing Position | Description |
|---|---|---|
| 1* | Upright | Remains upright for at least 30 minutes, with roughly 1/3 of the field of view occupied by benthos and 2/3 open water |
| 2* | Upright | Remains upright for at least 30 minutes, primarily looking at benthos or open water |
| 3 | Upright | Before 30 minutes the camera tips over or runs out of battery |
| 4 | Immediately Tips Over | Looking directly at the benthos or open water for entire duration of video |

* Indicates usability scores that were used in analysis

Table S3. The number of usable BRUV surveys conducted within each Marine Protected area (MPA) and paired Reference Site (REF) from 2019 to 2022.

| Site | MPA Status | Number of Surveys |
|---|---|---|
| Anacapa Island | MPA | 157 |
| | REF | 109 |
| Carrington Pt. | MPA | 112 |
| | REF | 115 |

Table S4. Number of usable BRUV surveys per habitat classification, grouped by site and level of protection.

| Site | MPA Status | Hard | Mixed Hard | Mixed Soft | Soft |
|---|---|---|---|---|---|
| Anacapa Island | MPA | 17 | 46 | 33 | 61 |
| | REF | 12 | 37 | 26 | 34 |
| Carrington Pt. | MPA | 48 | 33 | 16 | 15 |
| | REF | 37 | 28 | 19 | 30 |

Table S5. Table showing the number of qualifying grid cells by depth bin within the marine protected area (MPA) and reference area (REF) at Anacapa and Carrington Pt. and the total survey area for each MPA and reference area.

| | | **Depth Bin (m)** | | | **Study Area ($km^2$)** |
|---|---|---|---|---|---|
| | | **30-50** | **50-70** | **70-100** | |
| Anacapa | MPA | 50 | 45 | 33 | 12.03 |
| | REF | 167 | 179 | 167 | 24.02 |
| Carrington Pt. | MPA | 342 | NA | NA | 11.73 |
| | REF | 322 | NA | NA | 13.68 |

Table S6. For each species observed in this study: the targeted status, the total number of individuals observed, the total number of BRUV surveys on which they were observed, and the percentage of BRUV surveys overall that they were observed.

| Common Name | Target Status | Total Individuals Observed | Number of Surveys Observed | Percentage (%) of Surveys Observed |
|---|---|---|---|---|
| Ocean Whitefish | Targeted | 5767 | 414 | 84 |
| California Sheephead | Targeted | 603 | 243 | 49 |
| Copper Rockfish | Targeted | 506 | 235 | 48 |
| Vermilion Rockfish | Targeted | 459 | 191 | 39 |
| Painted Greenling | Non-targeted | 172 | 137 | 28 |
| Blue Rockfish | Targeted | 1760 | 134 | 27 |
| Halfbanded Rockfish | Non-targeted | 1336 | 125 | 25 |
| Rosy Rockfish | Targeted | 363 | 112 | 23 |
| Blackeye Goby | Non-targeted | 225 | 100 | 20 |
| Blacksmith | Non-targeted | 1291 | 98 | 20 |
| Lingcod | Targeted | 117 | 93 | 19 |
| Olive Yellowtail Rockfish complex | Targeted | 168 | 90 | 18 |
| Pacific Sanddab | Targeted | 440 | 74 | 15 |
| Gopher Rockfish | Targeted | 100 | 63 | 13 |
| Giant Sea Bass | Non-targeted | 50 | 46 | 9 |
| Treefish | Targeted | 56 | 43 | 9 |
| Widow Rockfish | Targeted | 171 | 34 | 7 |
| Flag Rockfish | Targeted | 41 | 29 | 6 |
| White Seaperch | Non-targeted | 70 | 28 | 6 |
| Pile Perch | Non-targeted | 102 | 28 | 6 |
| Bocaccio | Targeted | 79 | 28 | 6 |
| Senorita | Non-targeted | 148 | 27 | 5 |
| Canary Rockfish | Targeted | 42 | 23 | 5 |
| Squarespot Rockfish | Targeted | 72 | 22 | 4 |
| Jack mackerel | Targeted | 1435 | 19 | 4 |
| Shortspine Longspine Combfish complex | Non-targeted | 20 | 16 | 3 |
| Kelp Bass | Targeted | 20 | 15 | 3 |
| Starry Rockfish | Targeted | 26 | 14 | 3 |
| Ronquil Spp | Non-targeted | 15 | 13 | 3 |
| California Spotted Scorpionfish | Targeted | 16 | 13 | 3 |
| California Yellowtail | Targeted | 91 | 12 | 2 |
| Spotfin Surfpearch | NA | 24 | 12 | 2 |
| Bat Ray | Non-targeted | 11 | 11 | 2 |

| Common Name | Target Status | Total Individuals Observed | Number of Surveys Observed | Percentage (%) of Surveys Observed |
|---|---|---|---|---|
| California Halibut | Targeted | 13 | 8 | 2 |
| Sharpnose Seaperch | Non-targeted | 8 | 7 | 1 |
| Rubberlip Seaperch | Non-targeted | 11 | 7 | 1 |
| Island kelpfish | Non-targeted | 9 | 6 | 1 |
| Rock Wrasse | Non-targeted | 5 | 5 | 1 |
| Pacific Sardine | Targeted | 284 | 5 | 1 |
| Lavender Scuplin | Non-targeted | 4 | 4 | 1 |
| Honeycomb Rockfish | Non-targeted | 5 | 4 | 1 |
| Californian Anchovy | Targeted | 1233 | 4 | 1 |
| Brown Rockfish | Targeted | 6 | 4 | 1 |
| Cabezon | Targeted | 4 | 4 | 1 |
| Kelp Rockfish | Targeted | 3 | 3 | 1 |
| Greenspotted Rockfish | Targeted | 5 | 3 | 1 |
| Calico Rockfish | Targeted | 3 | 3 | 1 |
| Sculpin Spp | Non-targeted | 3 | 2 | <1 |
| Opaleye | Non-targeted | 2 | 2 | <1 |
| Halfmoon | Non-targeted | 4 | 2 | <1 |
| Ocean Sunfish | Non-targeted | 2 | 2 | <1 |
| Pink Seaperch | Non-targeted | 2 | 2 | <1 |
| Topsmelt | NA | 70 | 2 | <1 |
| Swell Shark | Non-targeted | 1 | 1 | <1 |
| Black Perch | Non-targeted | 1 | 1 | <1 |
| Garibaldi | Non-targeted | 2 | 1 | <1 |
| Bluebanded Goby | Non-targeted | 2 | 1 | <1 |
| C-O turbot | Non-targeted | 1 | 1 | <1 |
| California Lizardfish | Non-targeted | 1 | 1 | <1 |
| Rainbow Seaperch | Targeted | 2 | 1 | <1 |
| Pacific Angel Shark | Targeted | 1 | 1 | <1 |
| Grass Rockfish | Targeted | 1 | 1 | <1 |
| Stripetail Rockfish | Targeted | 1 | 1 | <1 |
| Soupfin Shark | NA | 1 | 1 | <1 |

Table S7. Model results from the negative binomial generalized linear model testing the effect of MPA status, habitat type, depth, and the interaction between habitat and MPA status on the MaxN and biomass of targeted rockfishes at Carrington Pt. and Anacapa. We report the estimate effect size, z-score, and resulting p-value. P-values less than 0.5 are bolded. We also report the deviance explained for each model.

| | | Anacapa | | | | | | Carrington Pt. | | | | | |
|---|---|---|---|---|---|---|---|---|---|---|---|---|---|
| | | MaxN | | | Biomass | | | MaxN | | | Biomass | | |
| | | Estimate | Z score | P-value | Estimate | Z score | P-value | Estimate | Z score | P-value | Estimate | Z score | P-value |
| **MPA Status** | | 0.76 | 1.5 | 0.13 | 0.19 | 0.3 | 0.75 | -0.34 | -1.2 | 0.23 | -0.58 | -2.1 | **0.04** |
| **Habitat** | **Mixed Hard** | -0.83 | -2.1 | **0.03** | -0.87 | -1.9 | 0.06 | -0.31 | -1.1 | 0.26 | -0.33 | -1.2 | 0.22 |
| | **Mixed Soft** | -1.60 | -3.8 | **<0.001** | -1.69 | -3.4 | **<0.001** | -1.25 | -3.4 | **<0.001** | -1.30 | -3.5 | **<0.001** |
| | **Soft** | -2.44 | -6.1 | **<0.001** | -1.55 | -3.4 | **<0.001** | -2.46 | -5.8 | **<0.001** | -2.25 | -5.0 | **<0.001** |
| **Depth** | | 0.05 | 7.7 | **<0.001** | 0.05 | 6.2 | **<0.001** | 0.02 | 1.3 | 0.17 | 0.04 | 2.5 | **0.01** |
| **MPA Status/Mixed Hard** | | 0.57 | 1.0 | 0.33 | 0.63 | 0.9 | 0.36 | -0.09 | -0.2 | 0.82 | 0.05 | 0.1 | 0.90 |
| **MPA Status/Mixed Soft** | | 0.57 | 0.9 | 0.36 | 0.75 | 1.0 | 0.31 | 0.77 | 1.5 | 0.13 | 0.31 | 0.6 | 0.55 |
| **MPA Status/Soft** | | -0.22 | -0.4 | 0.72 | -1.43 | -1.9 | 0.06 | 0.39 | 0.8 | 0.46 | 0.83 | 1.5 | 0.13 |
| ***Deviance Explained*** | | | | *0.37* | | | *0.22* | | | *0.21* | | | *0.17* |

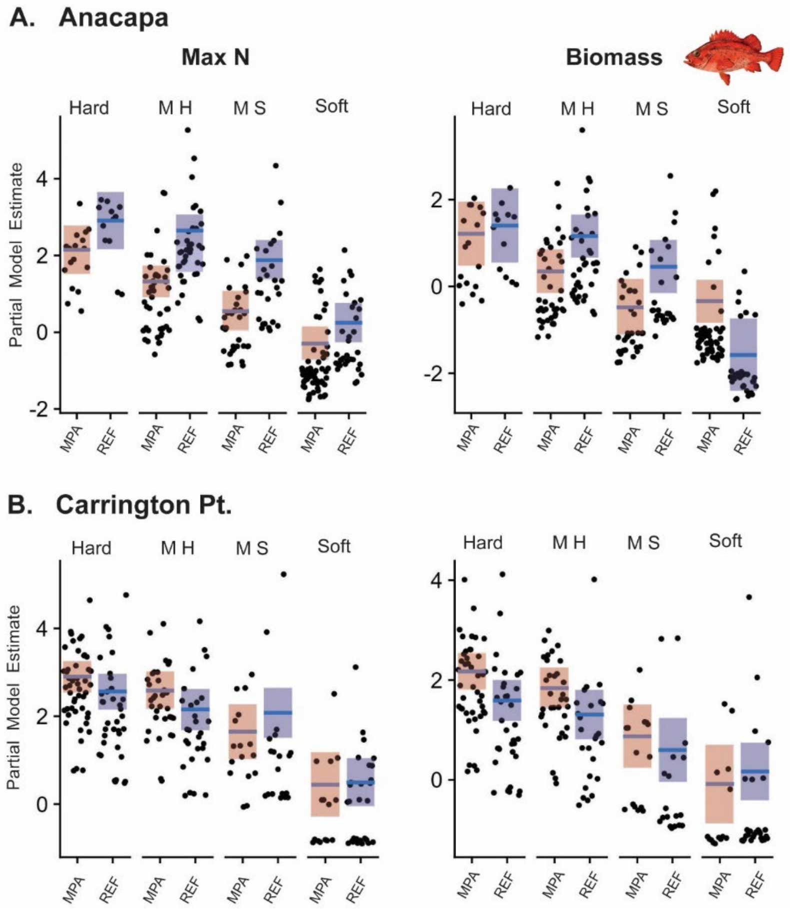


Figure S3. Partial model predictions from interaction term of MPA status and habitat from the negative binomial generalized linear model on the MaxN and biomass of targeted rockfishes. Panel A) shows the model results from Anacapa and panel B) shows the model results from Carrington Point. The blue line and shaded region represent the model predictions and 95% CI for a particular predictor variable when all other variables are held constant. The points represent the partial residuals of the model. To standardize the display across islands and metrics we set all predictions to 'MPA' and 'Hard' for MPA status and habitat, while depth was set to the median value for that particular data set. The habitat codes 'M H' and 'M S' represent mixed hard and mixed soft habitats respectively.

Table S8. Model results from the negative binomial generalized linear model testing the effect of MPA status, habitat type, depth, and the interaction between habitat and MPA status on the MaxN and biomass of ocean whitefish at Carrington Pt. and Anacapa. We report the estimate effect size, z-score, and resulting p-value. We also report the deviance explained for each model.

| | | Anacapa | | | | | | Carrington Pt. | | | | | |
|---|---|---|---|---|---|---|---|---|---|---|---|---|---|
| | | MaxN | | | Biomass | | | MaxN | | | Biomass | | |
| | | Estimate | Z score | P-value | Estimate | Z score | P-value | Estimate | Z score | P-value | Estimate | Z score | P-value |
| **MPA Status** | | -1.07 | -2.35 | **0.02** | -1.22 | 2.72 | **0.01** | -0.37 | -1.51 | 0.13 | -0.42 | -1.57 | 0.11 |
| Habitat | **Mixed Hard** | 0.36 | 1.51 | 0.13 | 0.57 | 1.82 | 0.07 | 0.36 | 1.51 | 0.13 | 0.31 | 1.23 | 0.22 |
| | **Mixed Soft** | -0.16 | -0.51 | 0.61 | 0.28 | 0.86 | 0.39 | -0.15 | -0.49 | 0.62 | -0.03 | -0.10 | 0.92 |
| | **Soft** | -0.68 | -1.96 | **0.05** | 0.04 | 0.11 | 0.91 | -0.67 | -1.94 | **0.05** | -0.73 | -1.96 | **0.05** |
| **Depth** | | -0.08 | -6.30 | **<0.001** | -0.03 | -6.52 | **<0.001** | -0.08 | -6.28 | **<0.001** | -0.06 | -4.70 | **<0.001** |
| **MPA Status/Mixed Hard** | | 0.01 | 0.02 | 0.98 | -0.18 | -0.35 | 0.73 | 0.01 | 0.04 | 0.96 | -0.15 | -0.38 | 0.70 |
| **MPA Status/Mixed Soft** | | 0.62 | 1.42 | 0.16 | 0.77 | 1.43 | 0.15 | 0.63 | 1.43 | 0.15 | 0.25 | 0.54 | 0.59 |
| **MPA Status/Soft** | | 0.74 | 1.67 | 0.10 | 0.11 | 0.21 | 0.84 | 0.74 | 1.68 | 0.09 | 0.36 | 0.74 | 0.46 |
| ***Deviance Explained*** | | | | *0.23* | | | *0.27* | | | *0.22* | | | *0.19* |

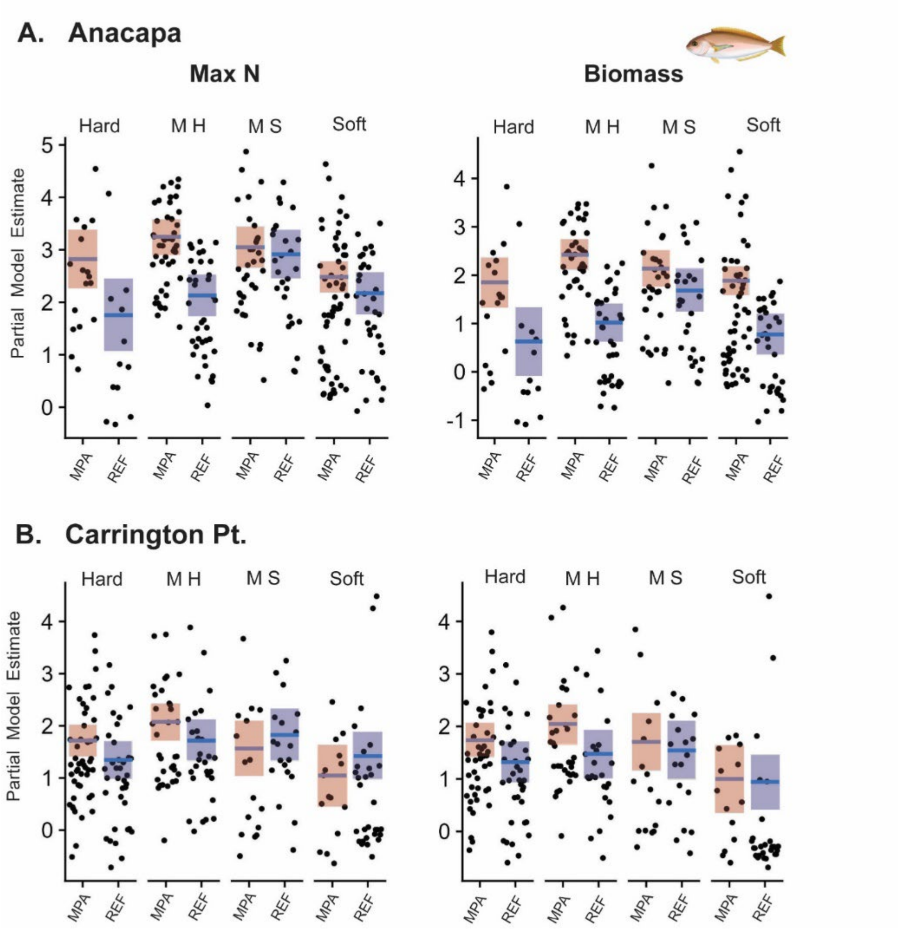


Figure S4. Partial model predictions from interaction term of MPA status and habitat from the negative binomial generalized linear model on the MaxN and biomass of ocean whitefish. Panel A) shows the model results from Anacapa and panel B) shows the model results from Carrington Point. The blue line and shaded region represent the model predictions and 95% CI for a particular predictor variable when all other variables are held constant. The points represent the partial residuals of the model. To standardize the display across islands and metrics we set all predictions to 'MPA' and 'Hard' for MPA status and habitat, while depth was set to the median value for that particular data set. The habitat codes 'M H' and 'M S' represent mixed hard and mixed soft habitats respectively.

Table S9. Model results from the negative binomial generalized linear model testing the effect of MPA status, habitat type, depth, and the interaction between habitat and MPA status on the MaxN and biomass of California Sheephead at Carrington Pt. and Anacapa. We report the estimate effect size, z-score, and resulting p-value. We also report the deviance explained for each model.

| | | Anacapa | | | | | | Carrington Pt. | | | | | |
|---|---|---|---|---|---|---|---|---|---|---|---|---|---|
| | | MaxN | | | Biomass | | | MaxN | | | Biomass | | |
| | | Estimate | Z score | P-value | Estimate | Z score | P-value | Estimate | Z score | P-value | Estimate | Z score | P-value |
| **MPA Status** | | 1.01 | 2.69 | **0.01** | 1.09 | 2.17 | **0.03** | -0.004 | -0.01 | 0.98 | -0.25 | -0.62 | 0.53 |
| Habitat | **Mixed Hard** | -0.14 | -0.43 | 0.67 | 0.53 | 1.27 | 0.20 | -0.01 | -0.05 | 0.96 | -0.18 | -0.47 | 0.63 |
| | **Mixed Soft** | -0.86 | -2.17 | **0.03** | -0.98 | -1.86 | 0.06 | -0.26 | -0.70 | 0.49 | -0.68 | -1.30 | 0.19 |
| | **Soft** | -1.13 | -3.09 | **<0.01** | -0.94 | -1.96 | **0.05** | -0.47 | -1.13 | 0.26 | -0.97 | -1.66 | 0.10 |
| **Depth** | | -0.06 | -9.13 | **<0.001** | -0.08 | -8.62 | **<0.001** | -0.05 | -3.37 | **<0.001** | -0.06 | -2.90 | **<0.01** |
| **MPA Status/Mixed Hard** | | 0.22 | 0.50 | 0.62 | -0.25 | -0.43 | 0.67 | -0.02 | -0.06 | 0.95 | -0.14 | -0.23 | 0.82 |
| **MPA Status/Mixed Soft** | | 0.58 | 1.11 | 0.27 | 0.52 | 0.74 | 0.46 | 0.19 | 0.38 | 0.70 | 0.26 | 0.36 | 0.72 |
| **MPA Status/Soft** | | -0.01 | -0.01 | 0.99 | -0.37 | -0.55 | 0.58 | -1.77 | -2.33 | **0.02** | -2.07 | -1.93 | **0.05** |
| ***Deviance Explained*** | | | | *0.46* | | | *0.45* | | | *0.19* | | | *0.21* |

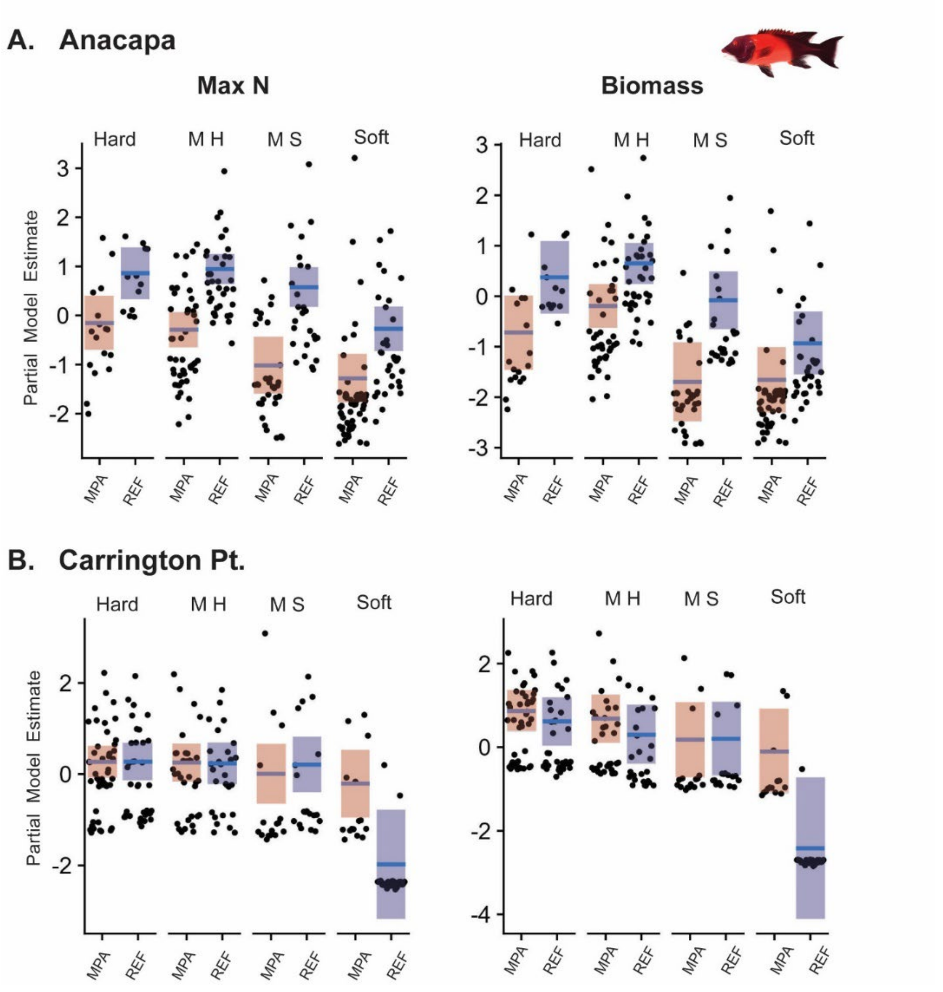


Figure S5. Partial model predictions from interaction term of MPA status and habitat from the negative binomial generalized linear model on the MaxN and biomass of California Sheephead. Panel A) shows the model results from Anacapa and panel B) shows the model results from Carrington Point. The blue line and shaded region represent the model predictions and 95% CI for a particular predictor variable when all other variables are held constant. The points represent the partial residuals of the model. To standardize the display across islands and metrics we set all predictions to 'MPA' and 'Hard' for MPA status and habitat, while depth was set to the median value for that particular data set. The habitat codes 'M H' and 'M S' represent mixed hard and mixed soft habitats respectively.

Methods S1.

Our BRUV stereo camera systems consist of a trapezoidal steel frame with two GoPro cameras (medium FOV, 30fps, 1080p) mounted in a calibrated stereo configuration. Each BRUV unit is equipped with two underwater dive lights to augment typically low ambient light conditions and a perforated PVC bait canister located in front of the camera array. To ensure accurate length measurements, we calibrated stereo camera systems before and after each sampling season (Harvey and Shortis 1998).

To standardize sampling location selection, we created a fishnet grid of 100m x 100m cells (ArcGIS Pro2.1) applied across each MPA and its respective reference area. Grid cells were allocated into three depth bins (30-50m, 50-70m, and 70-100m), and habitat maps from the California Seafloor Mapping Program (Golden and Cochrane 2013) were used to calculate the amount of hard bottom within each grid cell. Grid cells that contained >15% hard bottom for the two shallowest depth bins and >5% hard bottom for the deepest depth bin were selected as eligible sampling locations. Deeper grid cells (70-100m) contained considerably less hard bottom and required the lower criteria to ensure an even number of potential sampling grid cells across the depth bins.

We selected daily BRUV sampling locations haphazardly from the pool of qualifying grids cells, making sure that surveys were set at a minimum distance of 250m between units to minimize the effects of bait plume and reduce the likelihood of fish being resampled. We baited BRUVs with whole, moderately scored mackerel (*Scomber japonicas*), which was replenished before each survey (Dorman et al. 2012; Jones et al. 2020), and deployed each BRUV for a minimum of 30 minutes (Harasti et al. 2015).

To compare the overall amount of hard bottom in our depth ranges between the MPA and reference areas we calculated the number of eligible grid cells within each particular depth bin (30-50m, 50-70m, and 70-100m). We also report the total study area of each MPA and reference area from which all grid cells, hard and soft, were analyzed. MPA area was calculated from defined MPA boundaries and, for the inshore boundary, the 30m isobath. Calculating the area of the reference zone was somewhat subjective because the eastern and western boundaries are not designated as they are for the MPA. At Anacapa, we used the 30m and 100m isobaths for the inshore and offshore boundary respectively, and for eastern and western boundaries we estimated the area along the reef contours. At Carrington Pt., we followed a similar protocol, however the northern/offshore boundary was set by the 50m isobath.
